# VCP acts downstream of tTAFs to downregulate mono-ubiquitinated H2A and promote spermatocyte differentiation in *Drosophila*

**DOI:** 10.1101/2022.12.21.521492

**Authors:** Tyler J. Butsch, Olga Dubuisson, Alyssa E. Johnson, K. Adam Bohnert

**Affiliations:** 202 Life Sciences Building, Department of Biological Sciences, Louisiana State University, Baton Rouge, LA USA 70803

**Keywords:** VCP, meiosis, spermatocyte, tTAF, Polycomb, spermatogenesis

## Abstract

Valosin-containing protein (VCP) binds and extracts ubiquitinated cargo to regulate protein homeostasis. While VCP has been studied primarily in aging and disease contexts, it also affects germline development. However, the precise molecular functions of VCP in the germline, particularly in males, are poorly understood. Using the *Drosophila* male germline as a model system, we find that VCP translocates from the cytosol to the nucleus as germ cells transition into the meiotic spermatocyte stage. Importantly, nuclear translocation of VCP appears to be one critical event stimulated by testis-specific TBP-associated factors (tTAFs) to drive spermatocyte differentiation. Like *tTAF* mutants, spermatocyte gene expression fails to properly activate in *VCP*-RNAi testes, and germ cells arrest in early meiosis. At a molecular level, VCP activity supports spermatocyte gene expression by downregulating a repressive histone modification, mono-ubiquitinated H2A (H2Aub), at this developmental transition. Remarkably, experimentally blocking H2Aub in *VCP*-RNAi testes is sufficient to overcome the meiotic-arrest phenotype and to promote development through meiosis. Collectively, our data highlight VCP as a novel downstream effector of tTAFs that downregulates H2Aub to facilitate meiotic progression.

**SUMMARY STATEMENT:** VCP promotes the downregulation of mono-ubiquitinated H2A (H2Aub), potentially by driving H2A turnover. VCP-dependent downregulation of H2Aub occurs downstream of testis-specific TBP-associated factors and supports spermatocyte gene expression and differentiation.

## INTRODUCTION

Valosin-containing protein (VCP) is a broadly expressed AAA+ ATPase that functions as a ubiquitin-selective protein segregase (Ye, et al., 2017). Given its essential activities in protein homeostasis, VCP has been studied extensively in relation to aging and degenerative diseases (Darwich, et al., 2020; Johnson, et al., 2015; Kakizuka, 2008; Laço, et al., 2012; Ritson, et al., 2010; Scarian, et al., 2022). However, VCP also acts in a variety of other biological contexts, including germ-cell development. In oocytes of the nematode *Caenorhabditis elegans*, CDC-48/VCP appears particularly significant during meiotic divisions, when it regulates chromosome condensation and segregation (Mouysset, et al., 2008; Sasagawa, et al., 2012; Sasagawa, et al., 2009; Sasagawa, et al., 2007). While there are indications that VCP may likewise perform regulatory roles during sperm development (Wu, et al., 2021), it is unclear at what stage VCP function may be most critical. Stage-specific transcriptomic data from *Drosophila* testes suggest that *ter94/VCP* is strongly expressed in meiotic-stage spermatocytes (Shi, et al., 2020). While this hints at a potentially important role for VCP at the spermatocyte stage, molecular functions of VCP in meiotic spermatocytes are unknown.

*Drosophila* spermatogenesis presents a genetically tractable model to investigate conserved mechanisms regulating male germ-cell development, as the steps involved in making a viable sperm cell are similar in flies and mammals (Griswold, 2016; Hennig, 1992). At the apical tip of the *Drosophila* testis, germline stem cells divide, producing a daughter cell that undergoes four rounds of mitotic division to yield 16 spermatogonia (Fig. 1A). Subsequently, spermatogonia differentiate into spermatocytes, which enter meiotic prophase (Fig. 1A). Extensive transcriptional rewiring occurs at meiotic prophase during spermatogenesis in flies (Fuller, 1998; White-Cooper, 2010), as in humans (Jan, et al., 2017). These changes to transcription promote meiotic progression and are also needed to support later stages in germ-cell development (Lin, et al., 1996; White-Cooper, et al., 1998). In *Drosophila*, testis-specific TBP-associated factors (tTAFs) are required to stimulate the spermatocyte gene expression program (Chen, et al., 2005; Chen, et al., 2011; Fuller, 1998; Hiller, et al., 2004; White-Cooper, et al., 1998). When tTAFs are non-functional, germ cells arrest as spermatocytes, causing male fly infertility (Lin, et al., 1996). Notably, a common form of human male infertility, non-obstructive azoospermia (NOA) with spermatocyte arrest, exhibits similar cellular and developmental defects (Meyer, et al., 1992; Soderström and Suominen, 1980) as *tTAF* mutants (Lin, et al., 1996). Identifying factors that support robust spermatocyte gene expression, perhaps in cooperation with tTAFs, could reveal fundamental controls relevant to the preservation of male fertility.

**Figure 1.**
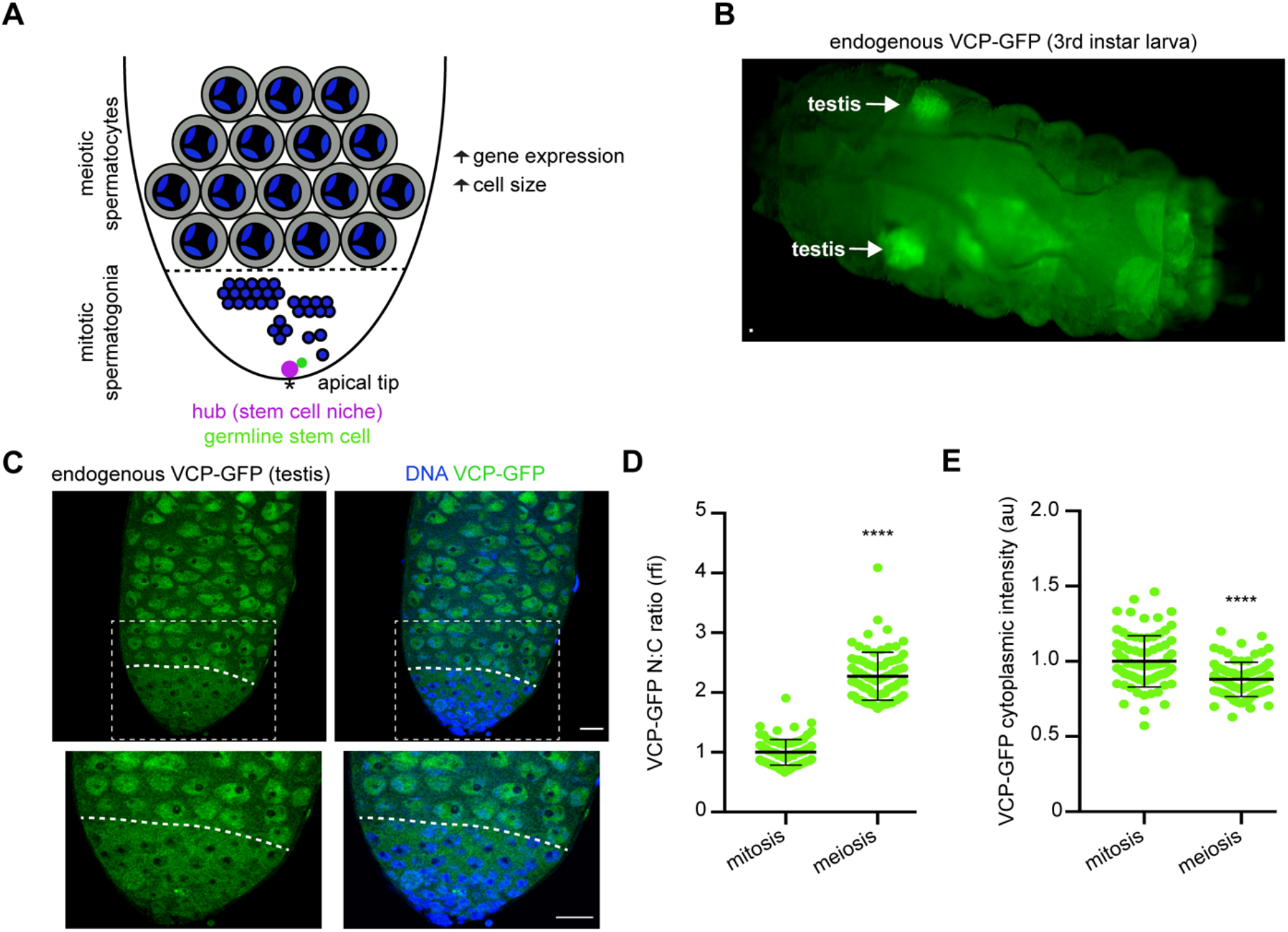
VCP expression is high in the *Drosophila* testis, particularly in spermatocyte nuclei. (**A**) Schematic of *Drosophila* spermatogenesis. The asterisk indicates the apical tip of the testis, where germline stem cells (green) and their niche (magenta) are located. Mitotic spermatogonia are indicated by solid blue circles, which correspond to chromatin. The dashed line indicates the mitotic-meiotic transition, when mitotic spermatogonia differentiate into meiotic spermatocytes (tri-lobed structures corresponding to paired bivalents in the nucleus). Spermatocytes exhibit significant cell growth and upregulated gene expression (right). (**B**) Image of a VCP-GFP 3^rd^ instar larva. The arrows indicate larval testes. (**C**) Images of Hoechst (DNA) and VCP-GFP in an adult testis. Images below are zoomed images of the outlined region in the top images. The dashed line indicates the mitotic-meiotic transition. (**D**) Quantification of the nuclear-cytoplasmic ratio (N:C) of VCP-GFP intensity in mitotic spermatogonia (*n*= 80 cells from 16 testes) and meiotic spermatocytes (*n*= 80 cells from 16 testes). Mean ± s.d. ****, p<0.0001. Wilcoxon matched pairs signed rank test. (**E**) Quantification of cytoplasmic VCP-GFP intensity in mitotic spermatogonia (*n*= 80 cells from 16 testes) and meiotic spermatocytes (*n*= 80 cells from 16 testes). Mean ± s.d. ****, p<0.0001. Paired t-test. Bars, 20 μm. See also Figure S1.

Molecularly, tTAFs have been postulated to drive spermatocyte gene expression by antagonizing the activity of a transcriptional repressor, Polycomb (Pc) (Chen, et al., 2005; Chen, et al., 2011). Pc is a core component of Polycomb Repressive Complex I (PRC1), which inhibits the expression of target genes, including meiosis-related genes (Endoh, et al., 2017; Zdzieblo, et al., 2014), by catalyzing H2A mono-ubiquitination (H2Aub) at repressed gene loci (Blackledge, et al., 2020; de Napoles, et al., 2004; Endoh, et al., 2008). Interestingly, downregulation of PRC1 activity is a requisite step for meiotic entry and cellular differentiation in several developmental contexts (Tang, et al., 2016; Yao, et al., 2018; Zdzieblo, et al., 2014). Despite these general themes, how downregulation of PRC1 activity is molecularly achieved during male germ-cell development remains incompletely understood, especially in relation to upstream signals from tTAFs. In principle, this may involve activation of factors that help to reverse repressive H2Aub epigenetic marks. However, the spatiotemporal regulation of H2Aub in the testis is unclear, as is the identity of enzymes that would facilitate its possible downregulation at meiosis.

In this study, we report that VCP operates downstream of tTAFs to downregulate H2Aub and to permit spermatocyte gene expression and development in the *Drosophila* testis. Interestingly, we find that VCP is cytoplasmic in mitotic spermatogonia but shuttles into the nucleus in meiotic spermatocytes. Nuclear translocation of VCP is dependent on tTAFs, and, like *tTAF* mutants, *VCP*-RNAi testes do not contain germ cells that develop beyond the spermatocyte stage. In line with these findings, we demonstrate that VCP is required to activate the expression of a subset of tTAF-regulated genes, and we observe that VCP translocation into the nucleus at meiotic entry coincides with a VCP-dependent downregulation of H2Aub. Remarkably, inhibiting PRC1 ubiquitin ligase activity is sufficient to advance sperm development past meiosis in the absence of VCP, suggesting H2Aub downregulation is an important function of VCP in spermatocyte differentiation. Overall, our findings identify VCP as an essential regulator of both spermatocyte gene expression and meiotic progression.

## RESULTS

### VCP is highly expressed in the *Drosophila* testis and enters the nucleus as germ cells progress into meiotic prophase

We recently generated a collection of fly strains to assess VCP function in multisystem proteinopathy (MSP-1) (Wall, et al., 2021). As part of these studies, we used CRISPR to knock-in a GFP tag at the endogenous *VCP* locus. While working with this strain, we noticed that VCP-GFP signal was exceptionally bright in 3^rd^ instar larval testes (Fig. 1B), which contain mostly spermatocytes (Gärtner, et al., 2014). Interestingly, previous RNA sequencing analyses have likewise suggested that VCP is highly expressed in the fly testis (Brown, et al., 2014; Leader, et al., 2018; Shi, et al., 2020). Thus, we sought to analyze VCP localization and regulatory dynamics during *Drosophila* spermatogenesis in more detail.

We imaged adult VCP-GFP testes such that we could observe endogenous VCP expression patterns in a fully developed male germline. While the protein was clearly expressed at multiple early stages in male germ-cell development, its localization varied dependent on developmental stage; VCP-GFP was primarily cytoplasmic in mitotic spermatogonia, but it translocated into the nucleus as cells developed into meiotic spermatocytes (Fig. 1C). Consequently, spermatocytes showed a higher nuclear-to-cytoplasmic VCP-GFP ratio compared to spermatogonia (Fig. 1D), and cytoplasmic VCP-GFP fluorescence was higher in spermatogonia than in spermatocytes (Fig. 1E). We confirmed that VCP enters the nucleus as germ cells exit mitosis and enter meiosis by imaging VCP-GFP in testes with germline-specific knockdown of *bag of marbles* (*bam*), which is required for mitotic spermatogonia to progress into the meiotic spermatocyte stage (McKearin and Spradling, 1990). Throughout *bam*-RNAi testes, VCP-GFP signal was restricted to the cytoplasm and undetectable in nuclei (Fig. S1), indicating that the nuclear translocation event occurs after the spermatogonial stage. These data support the conclusion that nuclear translocation of VCP is entrained with entry into meiotic prophase during spermatogenesis.

### Germline-specific knockdown of *VCP* arrests cells in meiotic prophase

Given the redistribution of VCP into the nucleus at meiotic prophase, we hypothesized that VCP may play an important role in post-mitotic stages of sperm development. To test this hypothesis, we knocked down *VCP* in the germline using a BamGal4 driver (Fig. S2A) and evaluated the developmental competency of male germ cells. As an initial assessment of whether mature sperm could even be produced in the absence of VCP, we first analyzed seminal vesicles. Not only were *VCP*-RNAi seminal vesicles noticeably shrunken, they were also devoid of mature sperm, as indicated by an absence of Hoechst-labeled needle-shaped sperm nuclei (Fig. 2A). Using DIC imaging, we confirmed that *VCP*-RNAi testes, which at a larger tissue level were also atypically small, lacked elongated spermatid bundles (Fig. 2B). Instead, the smaller *VCP-*RNAi testes appeared to show germ cells arrested at an earlier stage of development alongside some degenerating post-mitotic germ-cell cysts (Fig. 2B; arrowheads). Consistent with a defect in mature sperm production, *VCP*-RNAi males were completely infertile (Fig. S2B).

**Figure 2.**
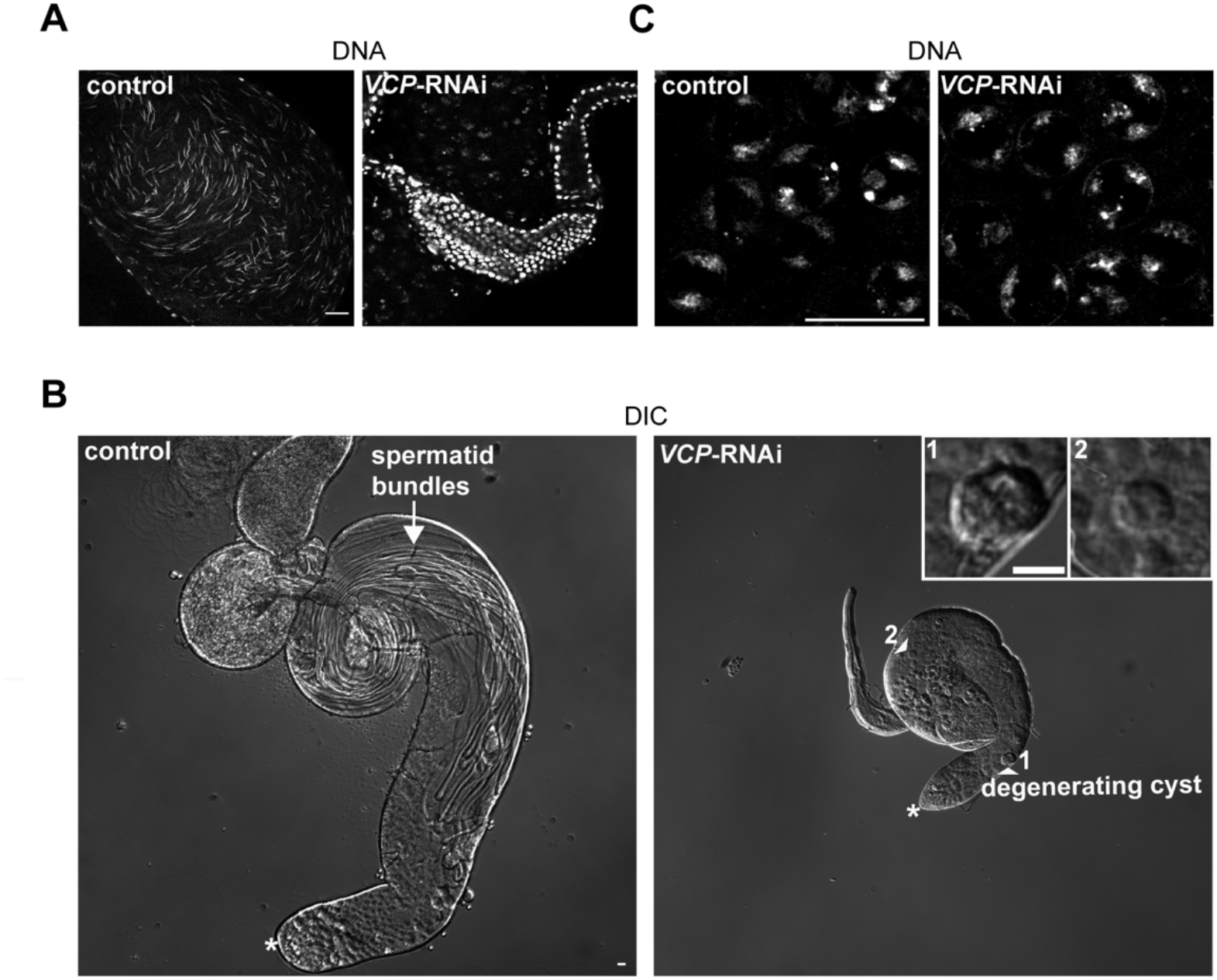
VCP is required for male fertility and meiotic progression. (**A**) Images of DNA (Hoechst) in control (BamGal4/+) and *VCP*-RNAi (BamGal4>*VCP*-RNAi) seminal vesicles. Mature sperm nuclei are needle-shaped (control), while only epithelial cells of the seminal vesicle are labeled in *VCP*-RNAi testes. (**B**) Low magnification DIC images showing whole control and *VCP*-RNAi testes. The arrow indicates elongated spermatid bundles in control testes. The arrowheads indicate degenerating germ-cell cysts in *VCP*-RNAi testes, which are also magnified in insets. Numbers next to each arrowhead correspond to each inset in the top right corner. The asterisks indicate the apical tip of each testis. (**C**) Images of Hoechst-labeled spermatocytes in control and *VCP*-RNAi testes. We did not observe cells that developed beyond this developmental stage in all *VCP*-RNAi testes analyzed. Bars, 20 μm. See also Figure S2.

To identify the precise developmental stage at which germ-cell development arrested in the absence of VCP, we labeled testes with Hoechst and imaged at high magnification. Notably, the most developed cells we observed in *VCP*-RNAi testes were spermatocytes with partially-condensed chromatin, which appeared morphologically-similar to control spermatocytes (Fig. 2C); we did not observe germ cells that developed beyond the spermatocyte stage in a total of 20 *VCP*-RNAi testes analyzed. Thus, VCP is required for male germ-cell development beyond the spermatocyte stage. This defect corresponds with the developmental timepoint at which VCP normally enters the nucleus, suggesting VCP likely carries out an essential cellular function inside the nucleus to support sperm development.

### VCP promotes spermatocyte gene expression downstream of tTAFs

Similar to VCP, tTAFs are required for progression beyond the spermatocyte stage (Lin, et al., 1996). Because loss of function in VCP and tTAFs causes a common developmental arrest at the spermatocyte stage, we hypothesized that VCP may function in the same pathway as tTAFs. Intriguingly, co-imaging of a tTAF protein, Spermatocyte arrest (Sa-GFP), with antibody-labeled VCP indicated that VCP enters spermatocyte nuclei shortly after Sa is expressed in spermatocytes (Fig. S3A). Based on this timing, we were curious whether tTAFs may promote the nuclear translocation of VCP in spermatocytes. Indeed, knockdown of *sa* and another tTAF, *meiosis I arrest* (*mia*), impeded nuclear translocation of VCP in spermatocytes (Fig. 3A, B). However, knockdown of *VCP* did not reciprocally affect Sa expression or localization (Fig. S3B), indicative of a unidirectional pathway. Thus, we concluded that VCP depends upon tTAFs to enter spermatocyte nuclei.

**Figure 3.**
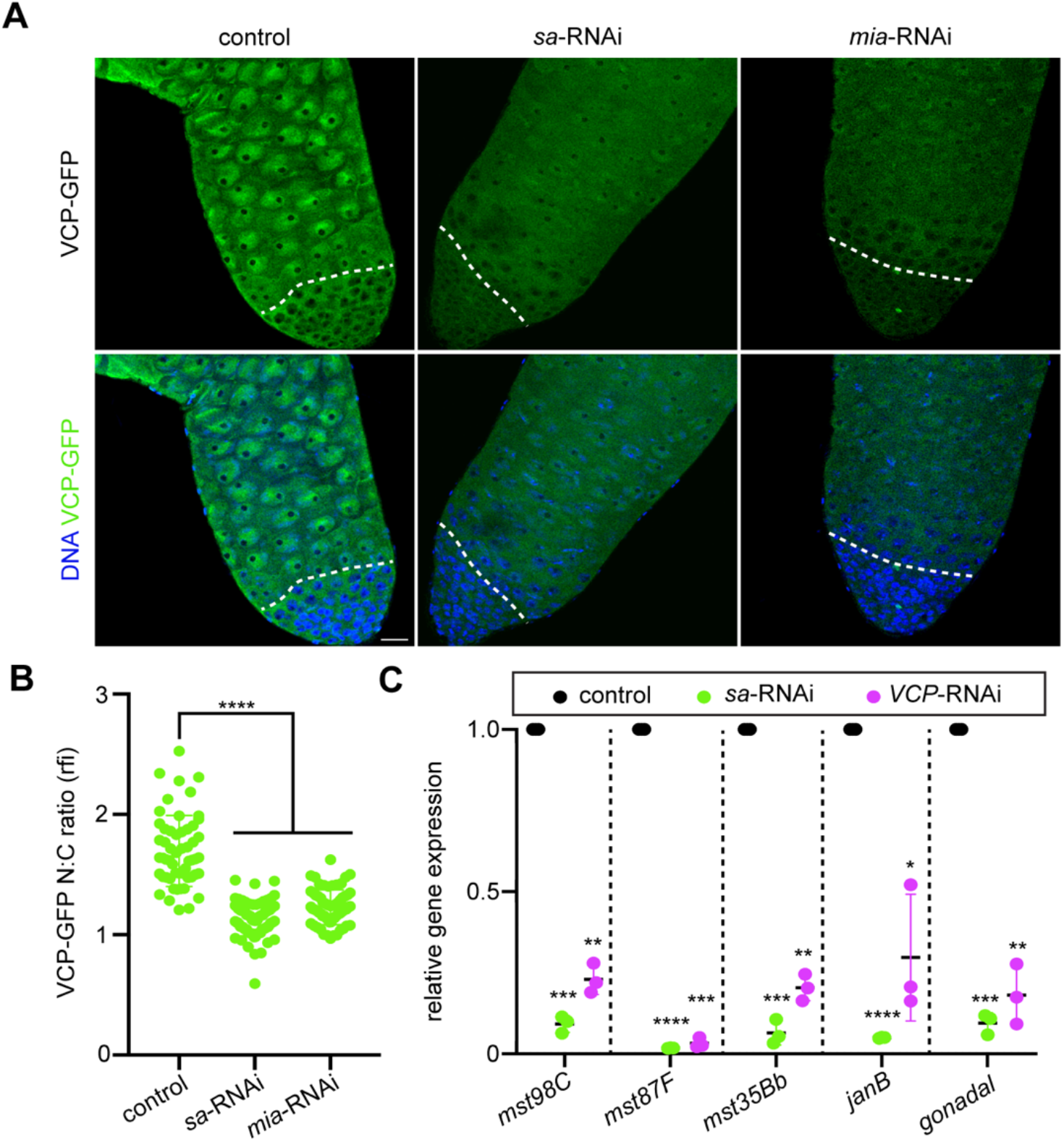
VCP enters spermatocyte nuclei downstream of tTAFs and drives spermatocyte gene expression. (**A**) Images of VCP-GFP and DNA (Hoechst) in control (BamGal4/+), *sa*-RNAi (BamGal4>*sa*-RNAi), and *mia*-RNAi (BamGal4>*mia*-RNAi) testes. The dashed lines indicate the mitotic-meiotic transition. Bar, 20 μm. (**B**) Quantification of the nuclear-cytoplasmic ratio (N:C) of VCP-GFP intensity in control (*n*= 55 spermatocytes from 11 testes), *sa*-RNAi (*n*= 90 spermatocytes from 18 testes), and *mia*-RNAi (*n*= 65 spermatocytes from 13 testes) testes. Mean ± s.d. ****, p<0.0001. Ordinary one-way ANOVA with Dunnett’s multiple comparisons test. (**C**) Graph of relative gene expression for the indicated tTAF-target genes. Gene expression was normalized to *actin* for each genotype (*n*= 3 replicates). *, p<0.05; **, p<0.01; ***, p<0.001; ****, p<0.0001. Two-way ANOVA. See also Figure S3.

A major function of tTAFs is to activate the spermatocyte gene expression program (Chen, et al., 2005; Chen, et al., 2011; White-Cooper, et al., 1998). Given that VCP enters the nucleus downstream of tTAFs and VCP also drives spermatocyte differentiation, we hypothesized that VCP may control the expression of tTAF-regulated genes. We tested this hypothesis by performing RT-qPCR to quantitatively measure the expression of a subset of tTAF-regulated genes (White-Cooper, et al., 1998) in control and *VCP-*RNAi testes. In agreement with our hypothesis, we consistently detected a sharp decrease in the expression of several tTAF-regulated genes *(mst98C, mst87F, mst35Bb, janB, gonadal)* in *VCP*-RNAi testes (Fig. 3C), potentially explaining the meiotic-arrest phenotype observed upon loss of VCP function.

### VCP and tTAFs stimulate H2Aub downregulation in spermatocytes

We next asked how VCP acts downstream of tTAFs to support target gene expression at the spermatocyte stage. tTAFs have been postulated to antagonize PRC1-mediated repression of meiotic gene expression in spermatocytes (Chen, et al., 2005; Chen, et al., 2011). We found that H2Aub, a repressive histone mark catalyzed by PRC1, decreased in wild type testes specifically at the spermatocyte stage; H2Aub was bright in spermatogonia and somatic cyst cells throughout the testis, but H2Aub signal declined dramatically in spermatocytes (Fig. S4A, B). In multiple contexts, VCP binds to and extracts ubiquitinated cargo (Ye, et al., 2017). As the reduction in H2Aub during spermatogenesis coincided with nuclear entry of VCP, we tested whether H2Aub downregulation in spermatocytes depended upon VCP function. Indeed, high H2Aub signal was retained in spermatocytes when *VCP* was knocked down (Fig. 4A), leading to an overall increase in H2Aub in protein extracts from whole testes (Fig. 4B, C). Knockdown of *sa* likewise showed an increase in H2Aub in spermatocytes (Fig. S5A) and protein extracts from whole testes (Fig. S5B, C), as would be expected if VCP operated downstream of tTAFs to trigger the downregulation of H2Aub.

**Figure 4.**
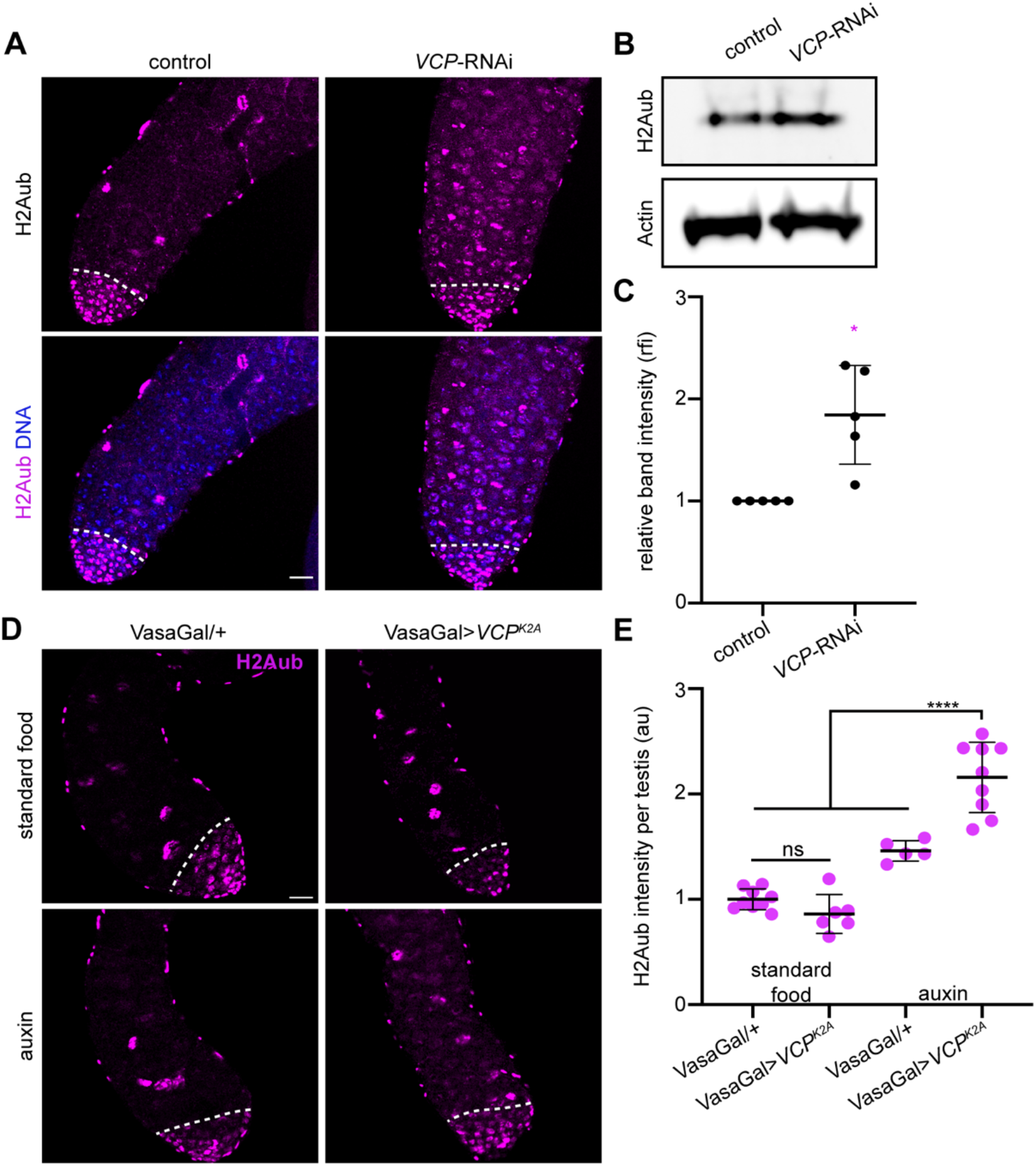
VCP negatively regulates H2A mono-ubiquitination. (**A**) Images of H2Aub and Hoechst (DNA) in control (BamGal/+) and *VCP*-RNAi (BamGal>*VCP*-RNAi) testes. The dashed lines indicate the mitotic-meiotic transition. (**B**) Western blotting for H2Aub (top) and Actin (bottom, loading control) in control and *VCP*-RNAi testes. (**C**) Quantification of H2Aub band intensity normalized to Actin band intensity (*n*= 5 replicates). Mean ± s.d. *, p<0.05. Unpaired t-test. (**D**) Images of H2Aub in VasaGal/+ and VasaGal> *VCP^K2A^* testes from flies that were fed standard food or auxin for 10 days. The dashed lines indicate the mitotic-meiotic transition. (**E**) Quantification of H2Aub intensity in spermatocytes of flies of the indicated genotypes fed the indicated types of food. Mean ± s.d. ns, p>0.05; ****, p<0.0001. Brown-Forsythe and Welch ANOVA. Bars, 20 μm. See also Figures S4 and S5.

To corroborate that H2Aub downregulation during spermatogenesis was dependent on VCP activity, we overexpressed a dominant allele of VCP that lacks ATPase activity (*VCP^K2A^*) (Chang, et al., 2021). To restrict transgene expression to adults, we utilized an auxin-inducible gene expression system (McClure, et al., 2022). This system utilizes a ubiquitously-expressed Gal4 inhibitor, Gal80, fused to an auxin inducible degron (AID). Upon auxin feeding, auxin binds Gal80 and causes its degradation, thus permitting Gal4 activity and transgene expression. For this experiment, we used the VasaGal4 driver, which is active in all germ cells (Demarco, et al., 2014) and should in principle provide the strongest transgene expression. Consistent with our *VCP*-RNAi data, H2Aub signal was higher than normal in spermatocytes of auxin-fed *VCP^K2A^* males relative to controls (Fig. 4D, E). Thus, we concluded that VCP ATPase activity is indeed required for H2Aub downregulation in spermatocytes.

### VCP supports H2A extraction and turnover

To gain insight into how VCP may downregulate H2Aub, we screened several known VCP cofactors for gene knockdowns that increased H2Aub in spermatocytes. Signficantly, we found that knockdown of *ufd1*, which encodes a ubiquitin-recognition factor that binds VCP and supports the sequestration and degradation of ubiquitinated proteins (Ramadan, et al., 2007; Sasagawa, et al., 2009; Shcherbik and Haines, 2007). Knockdown of *ufd1* blocked H2Aub downregulation in spermatocytes (Fig. 5A), suggesting that VCP may cooperate with Ufd1 to drive H2Aub downregulation.

**Figure 5.**
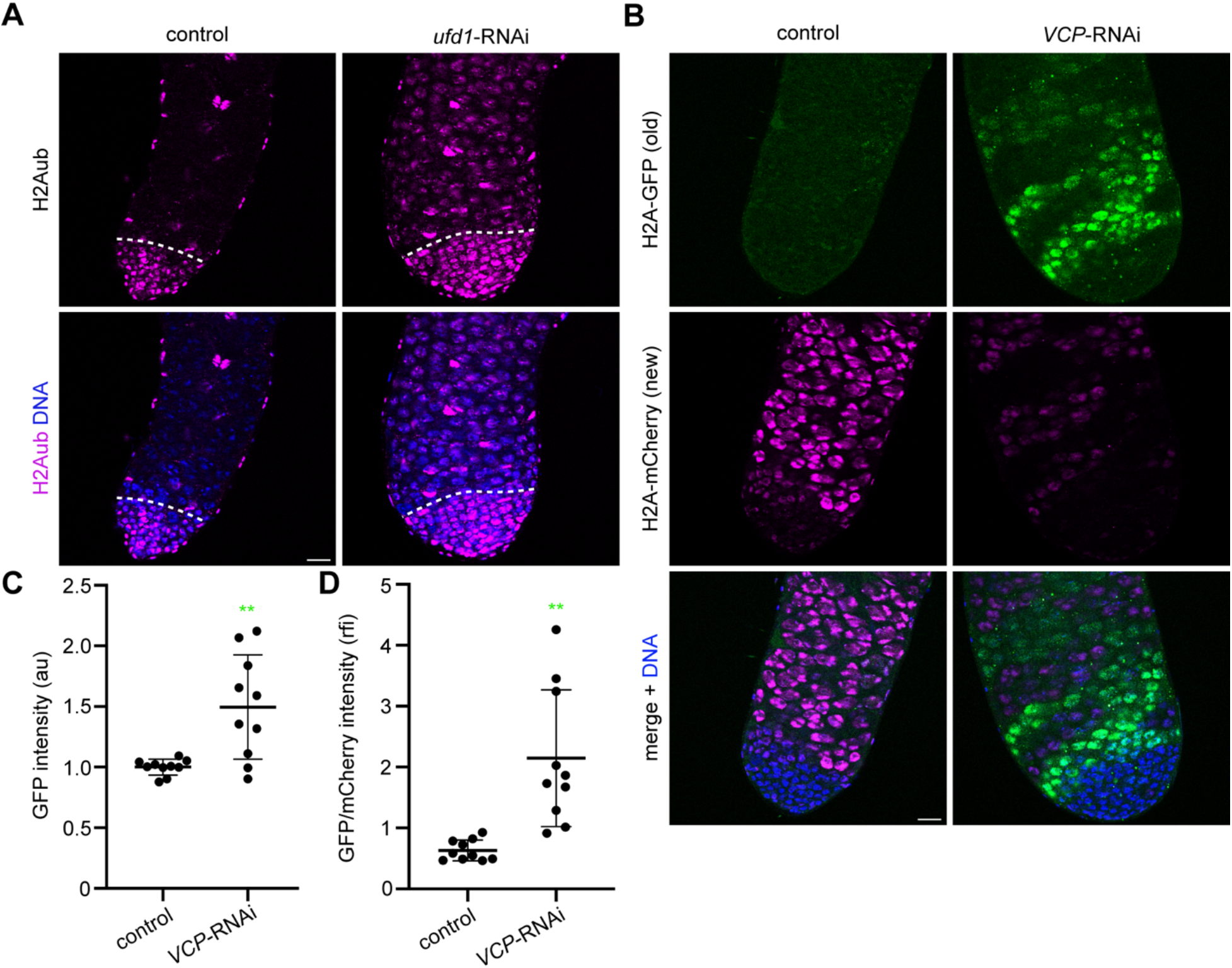
VCP cooperates with cofactor Ufd1 and promotes H2A turnover. (**A**) Images of H2Aub and Hoechst (DNA) in control (BamGal/+) and *ufd1*-RNAi (BamGal>*ufd7*-RNAi) testes. The dashed line indicates the mitotic-meiotic transition. (**B**) Images of Hoechst (DNA), H2A-GFP (old H2A), and H2A-mCherry (new H2A) in control (BamGal/+) and *VCP*-RNAi (BamGal>*VCP*-RNAi) testes. Testes were dissected and imaged five days after FLPase induction (H2A-GFP cassette removal). (**C**) Quantification of GFP intensity in spermatocytes of control (BamGal/+) and *VCP*-RNAi (BamGal> *VCP*-RNAi) testes. Mean ± s.d. **, p<0.01; Welch’s t-test. (**D**) Quantification of the GFP/mCherry ratio in spermatocytes of control (BamGal/+) and *VCP*-RNAi (BamGal> *VCP*-RNAi) testes (*n*= 10 testes for both genotypes in both graphs). Mean ± s.d. **, p<0.01; Welch’s t-test. Bars, 20 μm. See also Figure S6.

Because VCP and Ufd1 commonly act together to drive the extraction and degradation of ubiquitinated substrates, we hypothesized that VCP and Ufd1 may stimulate H2Aub downregulation by promoting the turnover of H2A in spermatocytes. To test this hypothesis, we employed a genetically-encoded H2A-turnover monitoring system (Kahney, et al., 2021). To activate this system, we used heat shock to transiently induce expression of a FLPase, which removes the H2A-GFP cassette and permits H2A-mCherry expression. Given time, older H2A will remain marked by GFP, whereas nascent H2A will be marked by mCherry. In control testes five days after heat shock, green signal was barely detectable above background levels whereas red signal was bright (Fig. 5B), consistent with high H2A turnover under normal conditions. We then asked if *VCP* depletion would cause a significant retention in GFP signal, as would be expected if VCP were required for H2A turnover. Indeed, five days after heat shock, GFP signal remained bright in many spermatocyte cysts of *VCP*-RNAi testes (Fig. 5B). Interestingly, cysts with bright GFP signal had almost no detectable mCherry signal; but mCherry signal was detectable, albeit still dimmer than controls, in cysts with dimmer GFP signal (Fig. 5B). Not only was the total GFP signal generally brighter in VCP-RNAi testes (Fig. 5C), but the GFP/mCherry ratio was also significantly higher in *VCP*-RNAi testes compared to controls (Fig. 5D). Thus, we concluded that VCP promotes the turnover of H2A in spermatocytes, which could potentially contribute to VCP-dependent H2Aub downregulation.

Ubiquitinated substrates extracted by VCP are often targeted to the proteasome for degradation (Ye, et al., 2017). Therefore, we next asked whether proteasomal degradation was also involved in H2Aub downregulation. Notably, mutation in a gene that codes for a proteasome component, *rpt2*, causes a meiotic arrest resembling *VCP* and *tTAF* mutants (Lu, et al., 2013). While germline-specific knockdown of *rpt2* in fact blocked downregulation of H2Aub in spermatocytes (Fig. S6A), VCP remained almost completely cytoplasmic in spermatocytes and failed to effectively translocate into the nucleus (Fig. S6B). Thus, while it remains possible that the proteasome may also act downstream of VCP to degrade extracted H2Aub, it also apparently acts upstream in the control of VCP nuclear translocation.

### Genetically inhibiting H2Aub promotes gene expression and suppresses the meiotic-arrest phenotype in *VCP*-RNAi testes

In several systems, downregulating H2Aub and/or PRC1 activity directly permits developmental gene expression and cellular differentiation (Blackledge, et al., 2020; de Napoles, et al., 2004; Endoh, et al., 2008; Endoh, et al., 2012; Endoh, et al., 2017; Gutiérrez, et al., 2012; Pengelly, et al., 2015; Tamburri, et al., 2020; Tang, et al., 2016; Yao, et al., 2018; Zdzieblo, et al., 2014). We therefore hypothesized that the developmental defects observed in *VCP*-RNAi testes may in part be due to failure to reduce H2Aub. To test this, we used a BamGal4 driver to simultaneously knock down *VCP* and *sex combs extra* (*sce*), the PRC1 E3 ubiquitin ligase that catalyzes H2Aub (de Napoles, et al., 2004). As expected, H2Aub levels were dampened in spermatocytes of *VCP*-RNAi *sce*-RNAi double-knockdown (DKD) testes, such that H2Aub patterns resembled that of control spermatocytes (Fig. S7A). We then assessed the effects on spermatocyte gene expression. Notably, each of the five tTAF-target genes we previously found to be downregulated in *VCP*-RNAi testes (Fig. 3C) increased in expression in DKD testes, to levels comparable to those seen in control testes (Fig. 6A). To broaden our analysis, we analyzed five additional, putative tTAF targets (Jiang, et al., 2018; White-Cooper, et al., 1998) and confirmed that all but one (*htrA2*) were indeed expressed significantly higher in DKD testes compared to *VCP*-RNAi testes (Fig. 6A). These data fit with the model that VCP generally permits the expression of several tTAF-regulated genes by downregulating H2Aub.

**Figure 6.**
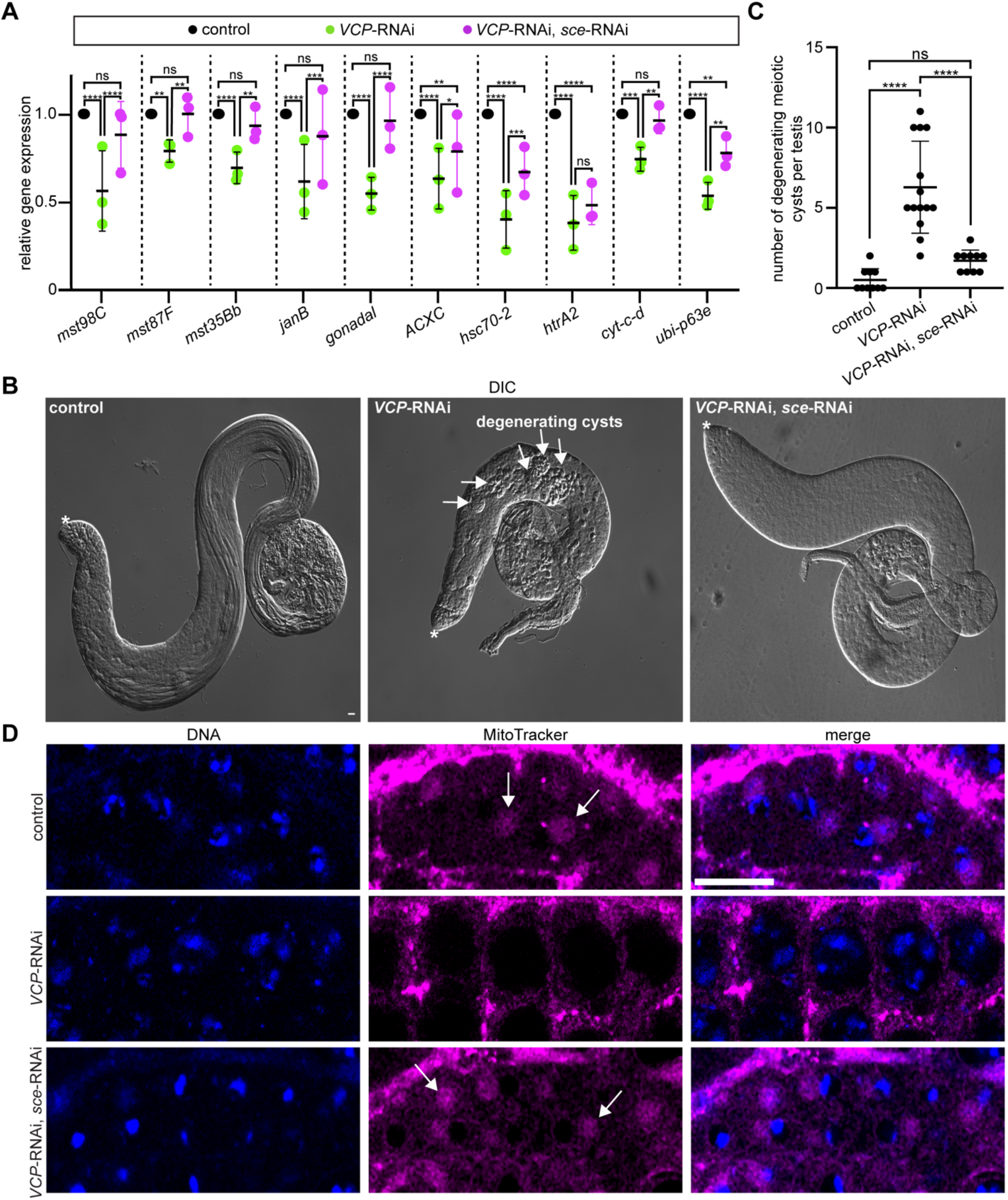
Inhibition of H2A mono-ubiquitination rescues meiotic arrest in the absence of VCP. (**A**) Graph of relative gene expression for the indicated tTAF-target genes. Gene expression was normalized to *actin* for each genotype (*n*= 3 replicates). Mean ± s.d. ns, p>0.05; *, p<0.05; **, p<0.01; ***, p<0.001; ****, p<0.0001. Two-way ANOVA. (**B**) Low magnification DIC images of control (BamGal/+), *VCP*-RNAi (BamGal>*VCP*-RNAi), and DKD testes (BamGal>*VCP*-RNAi, *sce*-RNAi). The asterisks indicate the apical tip of each testis. The arrows indicate degenerating germ-cell cysts in *VCP*-RNAi testes. (**C**) Graph of the number of degenerating germ-cell cysts per testis in control (*n*= 10 testes), *VCP*-RNAi (*n*= 14 testes), and DKD testes (*n*= 10 testes). Mean ± s.d. ****, p<0.0001; ns, p>0.05. Ordinary one-way ANOVA. (**D**) Images of MitoTracker and Hoechst (DNA) in control (BamGal/+), *VCP*-RNAi (BamGal>*VCP*-RNAi), and DKD (BamGal>*VCP*-RNAi, *sce*-RNAi) testes. Arrows indicate example nebenkerns, markers of the round spermatid stage only present in post-meiotic germ cells. Bars, 20 μm. See also Figures S7 and S8.

Like *VCP* and *sa*, multiple of the de-repressed genes, including *hsc70-2 and ubi-p63e*, are required for meiotic progression (Azuma, et al., 2021; Lu, et al., 2013). We therefore hypothesized that genetically inhibiting H2Aub may rescue some aspects of germ-cell development in *VCP*-RNAi testes. To test this hypothesis, we used microscopy to assess how far germ-cell development progressed in DKD testes. Although DKD testes still failed to produce elongated spermatid bundles or mature sperm (Fig. 6B), DKD testes were visibly larger than *VCP*-RNAi testes (Fig. 6B) and exhibited a significant reduction in degenerating post-mitotic germ-cell cysts, which were prevalent in *VCP*-RNAi testes (Fig. 6B, C). We hypothesized that this increased testis size and suppression of cyst degeneration may be indicative of the presence of more developed germ cells in DKD testes. Indeed, combined knockdown of *VCP* and *sce* allowed germ cells to develop beyond the spermatocyte stage to the round spermatid stage, indicated by compact chromatin morphology and accumulation of mitochondria in a nebenkern structure (Fabian and Brill, 2012) adjacent to germ-cell nuclei (Fig. 6D). Interestingly, double-knockdown of *sa* and *sce* did not similarly permit meiotic progression (Fig. S7B), suggesting that tTAFs likely perform some additional function independent of H2Aub downregulation that is needed to drive meiotic progression.

As a complementary approach to assess the potential rescuing effects of *sce* inhibition, we used mosaic analysis with a repressible cell marker (MARCM) to examine the effects of an *sce^KO^* allele (Gutiérrez, et al., 2012) in the germline. In this system (Lee and Luo, 1999), FLP-based recombination produces cells that are homozygous for the allele of interest (i.e., *sce^KO^*) and do not express Gal80, a Gal4 inhibitor (Duffy, 2002); uninhibited Gal4, in turn, drives expression of a GFP marker that enables the identification of MARCM clones (Lee and Luo, 1999). In contrast, GFP-negative cells in the same tissue, which possess at least one functional copy of the gene of interest (i.e., *sce*), express Gal80 and block Gal4 activity. With this set-up, we first ensured that GFP-positive *sce^KO^* homozygotes could not mono-ubiquitinate H2A by imaging H2Aub in spermatogonia two days-post clone induction (dpci). As expected, GFP-positive *sce^KO^* clones were negative for H2Aub, but neighboring GFP-negative clones were positive for H2Aub (Fig. S8A). Thus, this MARCM strategy provides a means to effectively block H2Aub cell-autonomously in developing germ cells.

Experimentally, the MARCM system can also be applied to drive RNAi knockdown of genes specifically in GFP-positive MARCM clones, but not in GFP-negative neighboring cells. In GFP-positive cells, Gal80 is absent, which permits Gal4 activity, and therefore expression of an RNAi transgene (i.e., *VCP*-RNAi). In GFP-negative cells, Gal80 is present, which inhibits Gal4 activity and RNAi transgene expression (Lee and Luo, 1999). We therefore used the MARCM system to generate germ-cell clones expressing *VCP*-RNAi that were also homozygous for a *sce^KO^* allele, or, as a control, a *sce^WT^* allele. We tested whether VCP was indeed knocked down in GFP-positive cells by labeling VCP with an antibody in MARCM testes. As expected, VCP was undetectable in GFP-positive germ-cell nuclei, though it remained detectable in neighboring GFP-negative germ cells (Fig. S8B). After confirming that this system functioned properly, we imaged testes at 7-9 dpci, when cells should be in post-meiotic stages (Chandley and Bateman, 1962). We observed GFP-positive round spermatids in 12/18 *VCP*-RNAi, *sce^KO^* MARCM testes (Fig. S8C). However, we never observed post-meiotic GFP-positive cysts in *VCP*-RNAi, *sce^WT^* MARCM testes (0/16 testes) (Fig. S8C). These data confirm that VCP downregulates H2Aub in spermatocytes to promote spermatocyte differentiation, though VCP also appears to execute additional functions that support further germline development past the spermatid stage.

## DISCUSSION

Although previous studies have demonstrated that VCP influences germ-cell health and development (León and McKearin, 1999; Sasagawa, et al., 2012; Sasagawa, et al., 2009; Sasagawa, et al., 2007; Wu, et al., 2021), fundamental questions remain regarding how VCP functions and is regulated in this biological context, particularly in the male germline. In this study, we found that VCP translocates from the cytosol to the nucleus at the mitotic-meiotic transition, and downregulates H2Aub to promote changes in gene expression needed for subsequent stages of spermatogenesis (Fig. 7). These findings present a new molecular framework for understanding developmental regulation of VCP and the mechanisms by which it ultimately controls male fertility.

**Figure 7.**
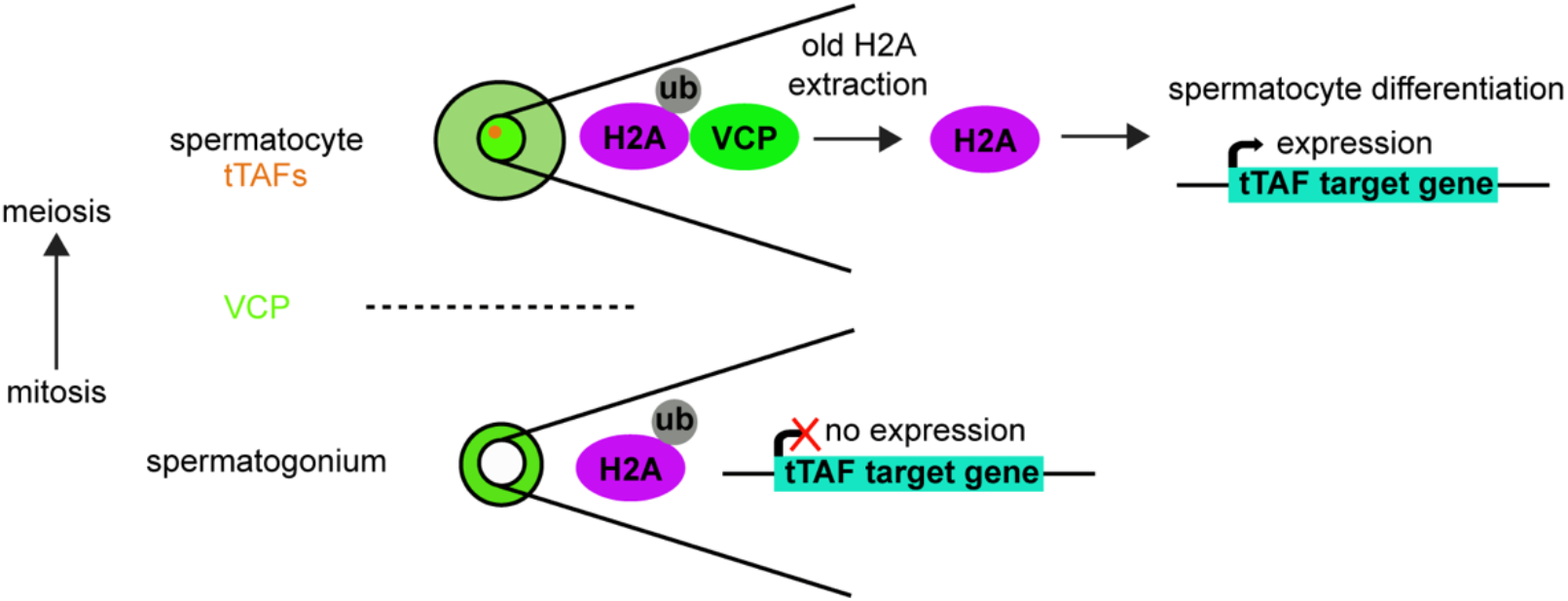
Model for VCP in spermatocyte development. In mitotic spermatogonia (below the dashed line), VCP (green) is cytosolic, H2Aub (H2A, magenta; ub, gray) is present, and tTAF-target genes (cyan) are not expressed. As spermatogonia differentiate into meiotic spermatocytes (above the dashed line), tTAFs (orange) are expressed and VCP enters spermatocyte nuclei. Nuclear entry of VCP triggers the extraction of old H2A and downregulation of H2Aub. The downregulation of H2Aub drives tTAF-target gene expression and spermatocyte differentiation.

In the *Drosophila* testis, nuclear entry of VCP at the spermatocyte stage acts as an initiating switch-like event that sets into motion a series of changes needed for developmental progression (Fig. 7). Molecularly, how the regulated nuclear entry of VCP occurs at this developmental stage is unknown; yet, it appears to be controlled at some level by upstream tTAFs. While tTAFs can bind and recruit transcriptional machinery important for the expression of spermatocyte and spermatid differentiation genes (Chen, et al., 2005; Chen, et al., 2011; Hiller, et al., 2004), direct binding between tTAFs and VCP has not been documented. Alternatively, other indirect methods of regulation may be at play. Structural studies have indicated that phosphorylation of VCP at a C-terminal tyrosine residue supports its nuclear translocation in other systems (Madeo, et al., 1998; Song, et al., 2015), raising the possibility that kinase signaling, perhaps downstream of tTAFs, could also potentially feed into this regulation. Notably, impairments to VCP nucleocytoplasmic shuttling are characteristic of degenerative pathologies, including MSP-1 (Song, et al., 2015), and it remains possible that similar factors may directly regulate VCP nuclear entry in somatic and germ cells. Thus, further clarifying controls on VCP nuclear translocation during spermatogenesis may not only identify entry points for treating forms of male infertility, including NOA, but may also reveal novel targets for treating VCP-related degenerative diseases.

Following nuclear entry, VCP then executes molecular functions important for spermatocyte gene expression. One important function of nuclear VCP appears to be downregulation of H2Aub, an activity that supports spermatocyte differentiation and the expression of at least some tTAF-regulated genes (Fig. 7). It is important to note that while VCP is essential for male germ-cell development, it may control only one branch of signaling downstream of tTAFs; whereas blocking H2Aub in the absence VCP permitted spermatocyte differentiation, blocking H2Aub in the absence of a tTAF, Sa, did not, as has also previously been shown (Chen, et al., 2005; Chen, et al., 2011). This suggests that tTAFs support spermatocyte differentiation through multiple signaling branches, which may be independently required. Indeed, tTAFs have been shown to also control other epigenetic modifications, including trimethylation of H3K4 (Chen, et al., 2005), hinting at possible branches besides VCP-H2Aub that may act in concert.

Mechanistically, our data suggest that VCP promotes H2Aub downregulation in spermatocytes via H2A extraction and turnover (Fig. 7). Because PRC1 catalyzes H2Aub at target gene loci (Blackledge, et al., 2020; de Napoles, et al., 2004; Endoh, et al., 2008), our findings support the model that PRC1 activity is indeed inhibitory to spermatocyte gene expression and differentiation, a point of contention in the spermatogenesis field (Chen, et al., 2005; Chen, et al., 2011; El-Sharnouby, et al., 2013). Interestingly, PRC1 components Pc and Sce are still present in spermatocytes (Chen, et al., 2005), where H2Aub levels are normally lowered. Thus, it is unclear how spermatocytes prevent mono-ubiquitination of new H2A in spermatocytes. Potentially, VCP may cooperate with an associated deubiquitinase, such as Asx and/or Calypso (ASXL and BAP1 in mammals, respectively) (Scheuermann, et al., 2010), or extract polyubiquitinated substrates involved in making this histone modification from target gene loci, similar to how it extracts Aurora B from chromatin in *Xenopus* embryos (Ramadan, et al., 2007). Consistent with the latter possibility, mutation of the major polyubiquitin gene, *ubi-p63E*, results in a similar meiotic arrest as *VCP*-RNAi (Lu, et al., 2013). Going forward, it will be important to clarify precisely how VCP downregulates H2Aub at the mitotic-meiotic transition, as this mechanism may also be pertinent to understanding the relief of PRC1 inhibition in other natural, developmental contexts.

## MATERIALS AND METHODS

### Fly husbandry and strains

Flies were maintained on standard cornmeal/agar food at 25°C, unless otherwise noted. For RNAi experiments, flies were incubated on standard cornmeal/agar food at 29°C for 5-7 days prior to dissection and imaging to boost Gal4 activity unless otherwise noted.

The following fly strains were used in this study: w^1118^, VCP-GFP (Wall, et al., 2021), UAS-*VCP*-RNAi (VDRC #24354), BamGal4 (Doug Harrison, Univ. Kentucky), *sa-GFP* (Chen, et al., 2005), UAS-*bam*-RNAi (BDSC #33631), UAS-*sa*-RNAi (BDSC #36730), UAS-*mia*-RNAi (BDSC #57790), UAS-*sce*-RNAi (BDSC #67924), *UAS-ufd1-* RNAi (BDSC #41823), UAS-*rpt2*-RNAi (BDSC #34795), *sce^KO^* (BDSC #80157), UAS-*GFP^nls^* (lab stock), VasaGal4; *hsFLP.D5*, UAS-*GFP*/cyo; *FRT82B*, tubGal80/TM6B (Butsch, et al., 2022), EyaGal4 (Leatherman and DiNardo, 2008), *UASp-FRT-H2A-eGFP-PolyA-FRT-H2A-mCherry* (Kahney, et al., 2021), tubGal80^AID^ (McClure, et al., 2022) (BDSC #92470), and *UAS-VCP^K2A^* (Chang, et al., 2021).

### Fertility assay

Single male flies were placed in a vial with 2-3 virgin females shortly after eclosion. Males were transferred to a fresh vial with new virgin females 2-3 days later. The presence or absence of progeny was scored after each mating. Males that failed to produce progeny in both matings were scored as infertile. Males that were able to produce progeny in both matings were scored as fertile. Flies were kept at 25°C on standard cornmeal/agar food for all matings.

### Larval imaging

3^rd^ instar larvae were washed several times in 1X phosphate-buffered saline (PBS; 137 mM NaCl, 2.7 mM KCl, 10 mM Na_2_HPO_4_, 1.8 mM KH_2_PO_4_) to remove food from larvae and prevent potential imaging artifacts. After washing, larvae were placed on a 4% agarose gel pad on a glass microscope slide and subsequently immobilized by firmly pressing a glass coverslip down on the larvae. Imaging was performed on a Leica M165 fluorescence stereomicroscope equipped with a GFP filter and 488 nm laser. Immunostaining, microscopy, and image processing

Testis immunohistochemistry was performed using standard procedures. Testes were dissected in 1X phosphate-buffered saline (PBS; 137 mM NaCl, 2.7 mM KCl, 10 mM Na_2_HPO_4_, 1.8 mM KH_2_PO_4_) and then immediately fixed in 4% paraformaldehyde for 20 minutes. Testes were washed three times in PBT (1X PBS, 0.1% Tween-20), then incubated in blocking buffer (3% BSA in 1X PBS) for at least 1 hour at room temperature. Testes were incubated with the primary antibody diluted in blocking buffer containing 2% Triton X-100 overnight at 4°C. The next day, testes were washed five times with PBT prior to applying the secondary antibody. Testes were incubated with the secondary antibody for at least 3 hours at room temperature in the dark. After the secondary antibody solution was removed, testes were washed five times with PBT. 1 μM Hoechst 33342 was added to the first wash to stain DNA. Testes were mounted in Vectashield antifade mounting medium prior to imaging.

The following antibodies were used in this study: rabbit anti-GFP (1:1000; Invitrogen A21311), mouse anti-GFP (1:200; Invitrogen A11120), rabbit anti-H2Aub (1:100; CST #8240S), rabbit anti-VCP (1:100; CST #2648), rabbit anti-mCherry (1:100; Invitrogen PA5-34975), goat anti-rabbit 488 (1:500; Invitrogen A11034), goat anti-rabbit Cy5 (1:500; Invitrogen A10523), goat anti-mouse 488 (1:500; Invitrogen A11001), goat anti-mouse Cy5 (1:500; Invitrogen A10524), and goat anti-rat 568 (1:500; Invitrogen A11077).

Images were acquired using an inverted Leica SP8 confocal microscope, equipped with a 40x objective (NA 1.30) and a white-light laser. Images were processed using Leica LAS X software, and quantifications were performed using Fiji (NIH) on 8-bit images prior to adjusting brightness or contrast.

### MitoTracker staining

Testes were dissected in PBT and immediately fixed in 4% paraformaldehyde for 20 minutes at room temperature. Testes were washed twice in PBT and then incubated at room temperature in 1:5000 MitoTracker Deep Red (ThermoFisher Cat# M22426) for 30 minutes. Testes were then washed twice in PBT and mounted in Vectashield antifade mounting medium prior to imaging.

### Germ-cell stage identification

Germ-cell staging was performed primarily based on chromatin morphology. Spermatocytes were identified based on chromatin features described in (Cenci, et al., 1994). Spermatids were identified based on chromatin and mitochondrial features described in (Fabian and Brill, 2012).

### Western blotting

Total protein was extracted from 40 pairs of testes per genotype. Testes were dissected in NP-40 lysis buffer containing protease inhibitors (6 mM Na_2_HPO_4_, 4 mM NaH_2_PO_4_, 1% NP40, 150 mM NaCl, 2 mM EDTA, 50 mM NaF, 0.1mM Na_3_VO_4_, 4 μg/ml leupeptin, one Roche cOmplete™ protease inhibitor tablet, pH 7.4). After dissections were complete, lysis buffer was removed, and testes were further lysed in 5x SDS Sample Buffer [1M Tris-HCl (pH 6.8), 1mM DTT, 20% SDS, 60% glycerol, Bromophenol Blue] using a sterile pestle. Samples were boiled at 100°C for 10 minutes, followed by a brief vortex and centrifugation for one minute. Proteins were resolved on a 4-12% Bis-Tris gel and transferred to a nitrocellulose membrane. Membranes were blocked in 4% milk for at least one hour. After blocking, membranes were incubated overnight with primary antibody at 4°C. The next day, membranes were washed three times for at least 10 minutes in PBT. Membranes were then incubated with secondary antibody for one hour at room temperature, followed by washing. Membranes were treated with an ECL chemiluminescent reagent (Bio-Rad) and proteins were visualized on a Bio-Rad ChemiDoc imaging system.

The following antibodies were used in this study: rabbit anti-H2Aub (1:2000; CST #8240S), mouse anti-actin (1:1000; Invitrogen MA5-11869), goat anti-rabbit IgG HRP (Invitrogen), and goat anti-mouse IgG HRP (Invitrogen).

### RT-qPCR

Total RNA was extracted from 50 pairs of testes per genotype by Trizol extraction. Total RNA was treated with DNase (Promega, M610A) to remove genomic DNA, followed by RNA precipitation with isopropanol and washes with 70% ethanol. 500 ng of RNA was used for cDNA synthesis using the iScript cDNA synthesis kit (Bio-Rad Cat# 1708891). qPCR was conducted using PowerUp™ SYBR^TM^ Green Master Mix (ThermoFisher). Data was analyzed using Design and Analysis software (v2.6). The 2^-ΔΔCt^ method was used to estimate the relative changes in gene expression. Primers used for each gene are below.

*actin*
F: TTGTCTGGGCAAGAGGATCAG
R: ACCACTCGCACTTGCACTTTC
*janB*
F: CTCCGTTTCAAAAATGTTACTCAACCG
R: CAACATCGGCGCCACG
*gdl*
F: GAACTTCCCGAAAACTTGCAGACAC
R: CGTTCTCCGCCTGCATCC
*mst35Bb*
F: GTGGAATGGCATAATTTCCATTTCTGC
R: GTTCACTGGTGGTGACCTTGC
*mst98C*
F: GCGGTCCTTGTAGTCCATGC
R: CTCCGGCACAATCTTCTCCG
*mst87f*
F: TCCGACTTGTCAAACCGATA
R: GCACGAAGGGTATCCACAAT
*ubi-p63E*
F: GGCTAAGATCCAAGACAAGGAA
R:GAGACGAAGGACCAAGTGAAG
*hsc70-2*
F: GAACCAGGTGGCCATGAAT
R: TCAGGTCCTCCTGTATCTTCTT
*cyt-c-d*
F: CCTCAAGGACCCGAAGAAATAC
R: TGACTTGAGGAAGGCAATCAA
*htrA2*
F: CTATCCGATGGCAGGACTTT
R: CCGAAAGATTGTTCACCTGTATG
*ACXC*
F: GGAACACAGTTACTTGAGGGAAA
R: CATCAGACGGACCTGGTAGTA

### MARCM

VasaGal4; *hsFLP.D5*, UAS-*GFP*/UAS-*VCP*-RNAi; *FRT82B, sce^KO^/FRT82B*, tubGal80 and VasaGal4; hs*FLP.D5*, UAS-*GFP*/UAS-*VCP*-RNAi; *FRT82B/FRT82B*, tubGal80 males were generated via standard crossing procedures (Butsch, et al., 2022). Males were heat shocked at 37°C the same day they eclosed for one hour to activate FLP expression and drive recombination. Following heat shock, flies were housed at 25°C on standard agar/cornmeal food until dissection. Testes were dissected and imaged at the indicated time-points following heat shock (“clone induction”).

### H2A turnover assay

*hsFLP.D5/+; BamGal4/UASp-FRT-H2A-eGFP-PolyA-FRT-H2A-mCherry* and hs*FLP.D5*/UAS-*VCP*-RNAi; *BamGal4/UASp-FRT-H2A-eGFP-PolyA-FRT-H2A-mCherry* males were generated via standard crossing procedures. Males were heat shocked at 37°C the same day they eclosed for one hour to activate FLP expression and drive removal of the *H2A-eGFP* cassette. Following heat shock, flies were housed at 25°C on standard agar/cornmeal food for five days until dissection, antibody labeling, and imaging.

### Auxin-induced temporal expression of VCP^K2A^

VasaGal4; tubGal80^AID^ flies were generated by standard crossing procedures and subsequently crossed to UAS-*VCP^K2A^* flies to generate VasaGal4; UAS-*VCP^K2A^*/+; tubGal80^AID^/+ males. Males were fed 5mM auxin or standard food (see above) at 29°C for 10 days prior to dissection and imaging. Flies were transferred to fresh food every 2-3 days.

### Statistical analyses

Information on sample size and statistics is provided in figure legends where applicable. Data normality was tested via the D’Agostino-Pearson test in combination with Q-Q plots prior to performing follow-up statistical analyses using GraphPad Prism software. Statistical tests used to determine significance are indicated in figure legends. The Student’s unpaired t-test was used when unpaired data for two groups were normally distributed and standard deviation was equal between both groups. Welch’s unpaired t-test was used when unpaired data for two groups were normally distributed, but standard deviation was not equal. The Mann-Whitney U-test was used when unpaired data for two groups were not normally distributed. The paired t-test was used when paired data for two groups were normally distributed and standard deviation was equal. The Wilcoxon matched pairs signed rank test was used when paired data for two groups were not normally distributed. The Brown-Forsythe ANOVA with Dunnett’s multiple comparisons test was used when there were more than two groups with normally distributed data, but standard deviation was not equal between the groups.

## ACKNOWLEDGEMENTS

We would like to thank Xin Chen (Johns Hopkins), Minx Fuller (Stanford), Doug Harrison (Univ. of Kentucky), and Tzu-Sang Kang (National Tsing Hua University, Taiwan) for generously providing fly strains needed to perform this study. We would also like to thank members of the Bohnert and Johnson labs at LSU and members of T.B.’s dissertation committee for providing helpful comments regarding the manuscript. Lastly, we would like to thank the LSU Shared Instrumentation Facility and LSU Genomics Facility for providing tools to complete this study.

## DATA AVAILABILITY

This study did not generate large datasets that were deposited in a repository. All data that were generated in this study can be provided by the corresponding author (K.A.B) upon request.

## Supplemental Figure Legends

**Figure S1.**
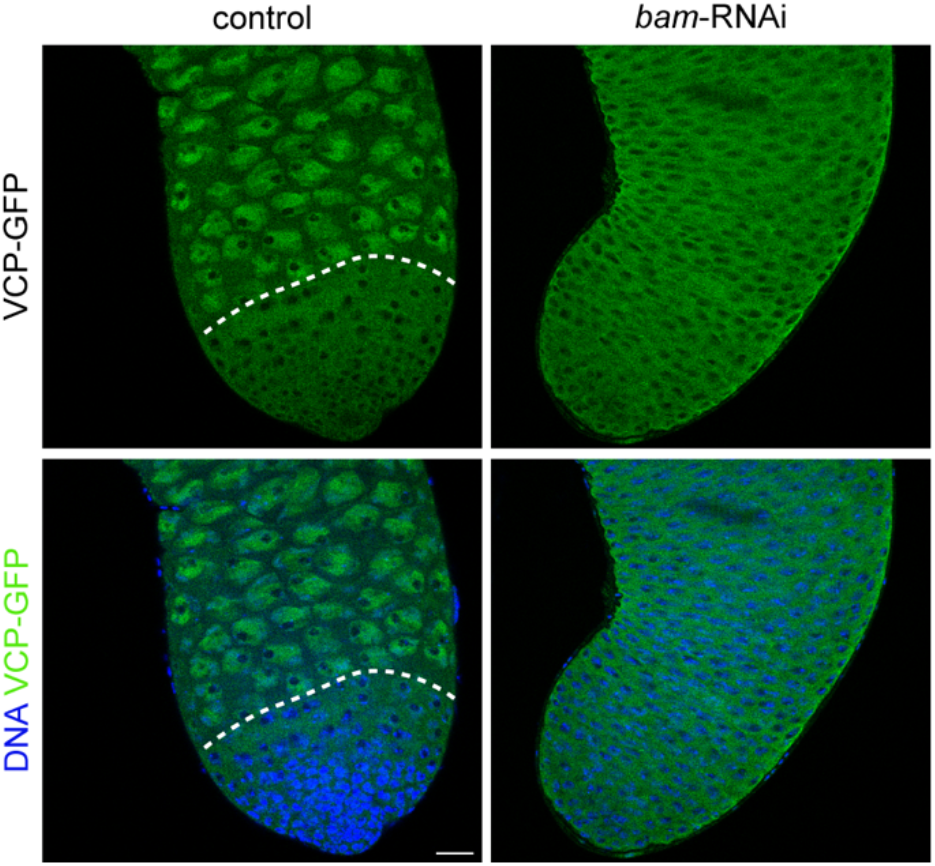
VCP is cytosolic in spermatogonia, but nuclear in spermatocytes. (**A**) Images of Hoechst (DNA) and VCP-GFP in control (BamGal/+) and *bam*-RNAi (BamGal>*bam*-RNAi) testes. The dashed line indicates the mitotic-meiotic transition. Bar, 20 μm.

**Figure S2.**
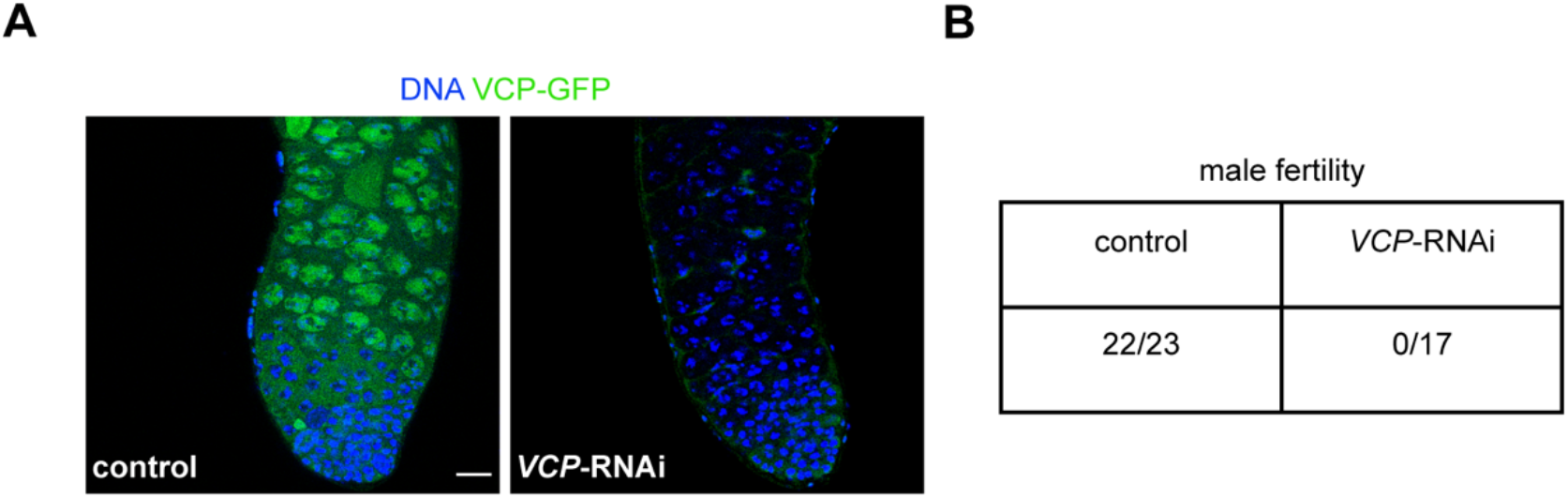
Germline-specific knockdown of VCP causes male infertility. (**A**) Images of Hoechst (DNA) and VCP-GFP in control (BamGal/+) and *VCP*-RNAi (BamGal> *VCP*-RNAi) testes. Bar, 20 μm. (**B**) Table of the proportion of males that are fertile.

**Figure S3.**
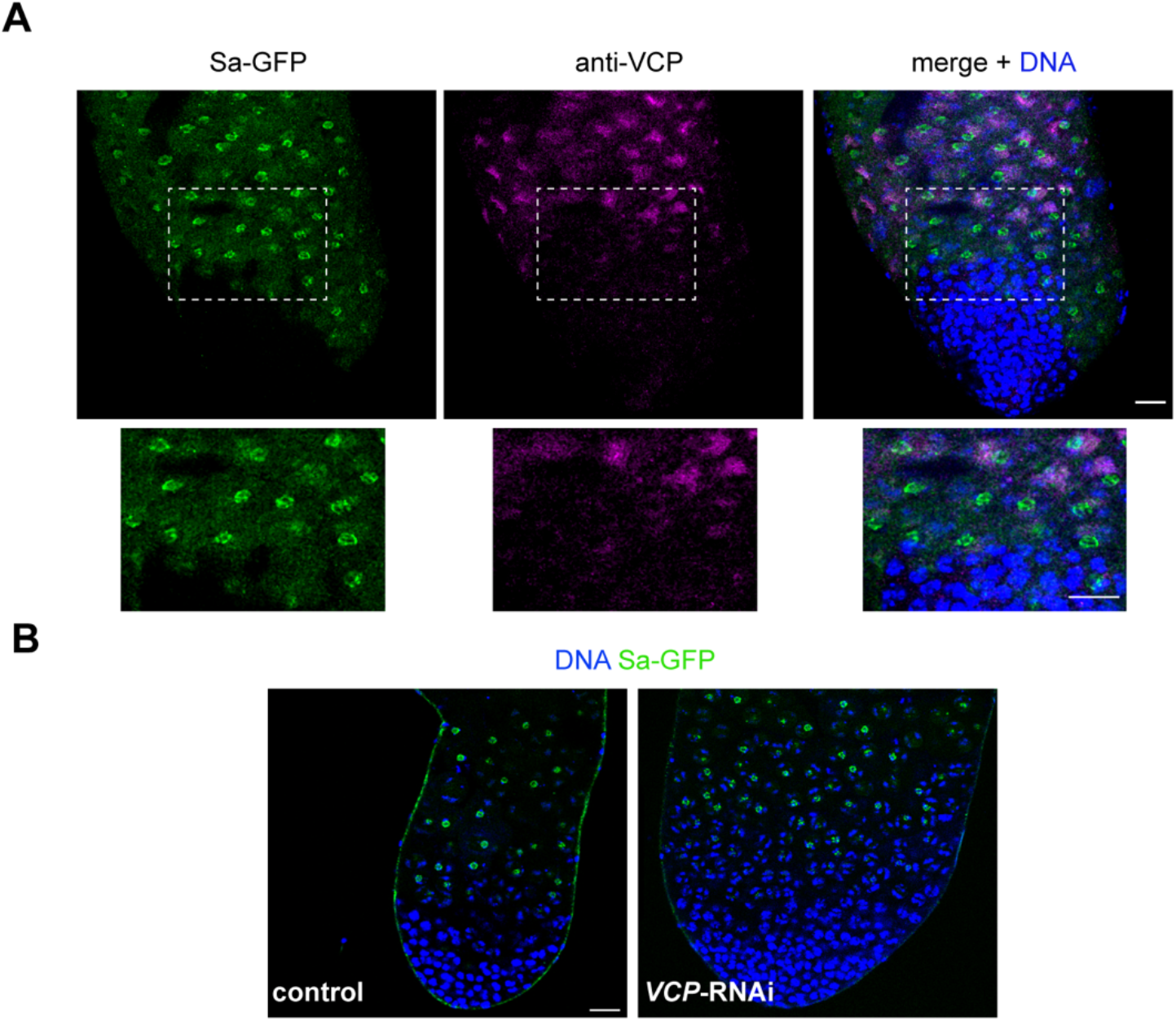
VCP acts downstream of tTAFs. (**A**) Images of Sa-GFP, VCP (anti-VCP), and Hoechst (DNA) in adult testes. Outlines indicate the region shown in the insets below. (**B**) Images of Sa-GFP and Hoechst (DNA) in control (BamGal/+) and *VCP*-RNAi (BamGal>*VCP*-RNAi) testes. Bars, 20 μm.

**Figure S4.**
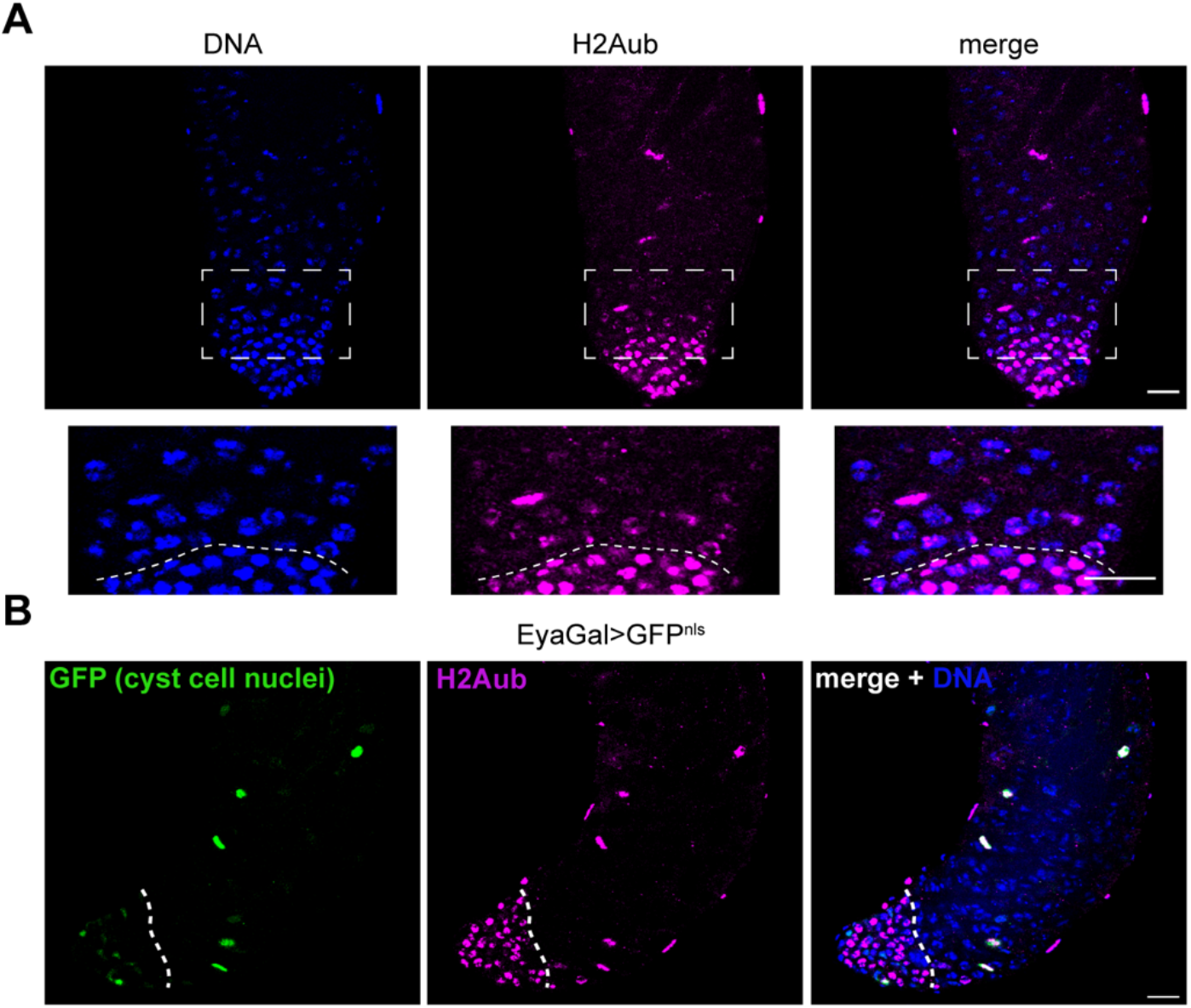
H2Aub is downregulated in germ cells at the mitotic-meiotic transition but not in somatic cyst cells. (**A**) Images of H2Aub and Hoechst (DNA) in wild type (*w^1118^*) testes. Outlines indicate the region shown in the insets below. The dashed line indicates the mitotic-meiotic transition. (**B**) Images of GFP (cyst cell nuclei), H2Aub, and Hoechst (DNA) in EyaGal>GFP^nls^ testes. The dashed line indicates the mitotic-meiotic transition. Bars, 20 μm.

**Figure S5.**
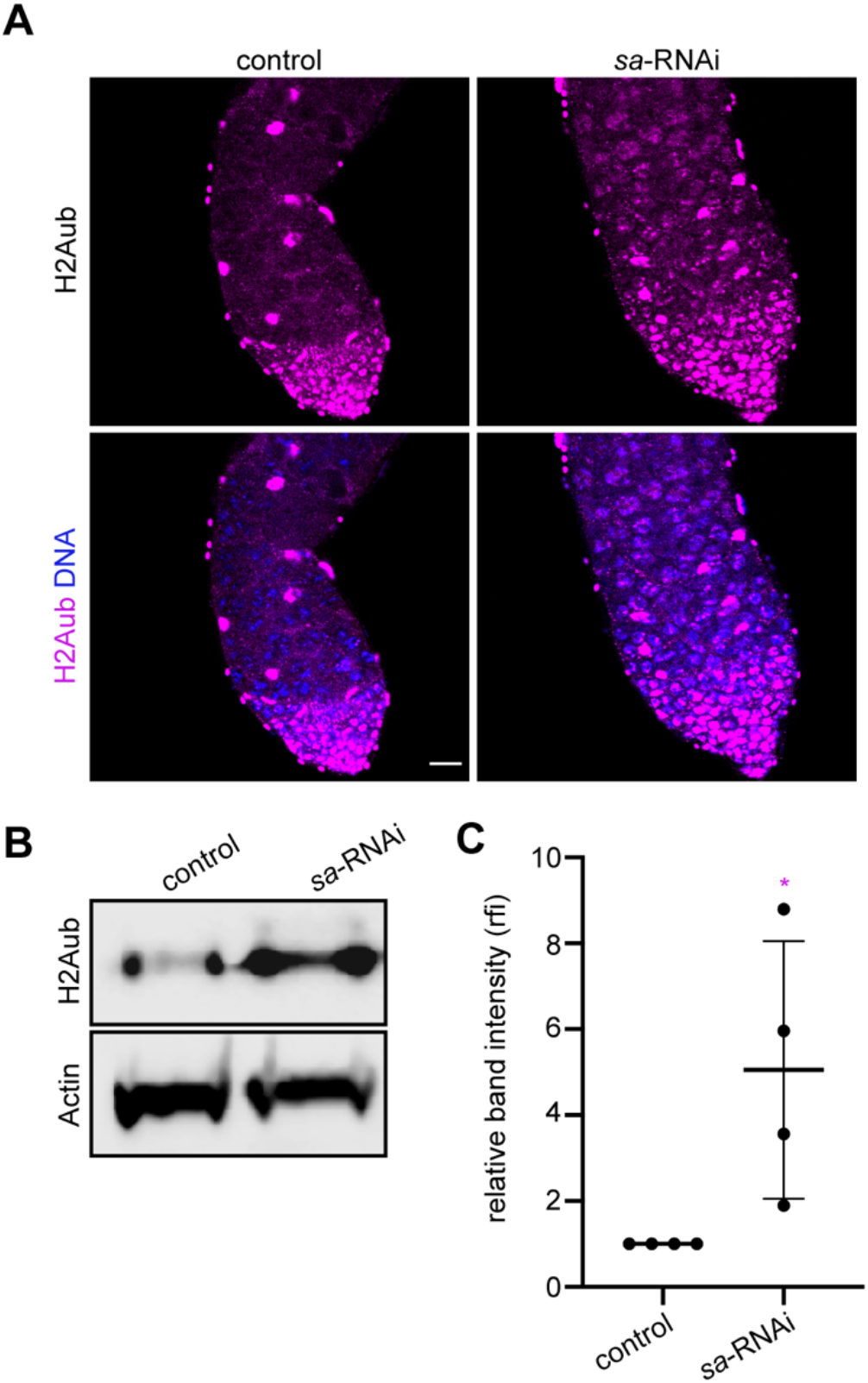
The tTAF, Sa, promotes H2Aub downregulation in spermatocytes. (**A**) Images of H2Aub and Hoechst (DNA) in control (BamGal4/+) and *sa*-RNAi (BamGal4>*sa*-RNAi) testes. Bar, 20 μm. (**B**) Western blotting for H2Aub (top) and Actin (bottom, loading control) in control and *sa*-RNAi testes. (**C**) Quantification of H2Aub band intensity normalized to Actin band intensity (*n*= 4 replicates). Mean ± s.d. *, p<0.05. Unpaired t-test.

**Figure S6.**
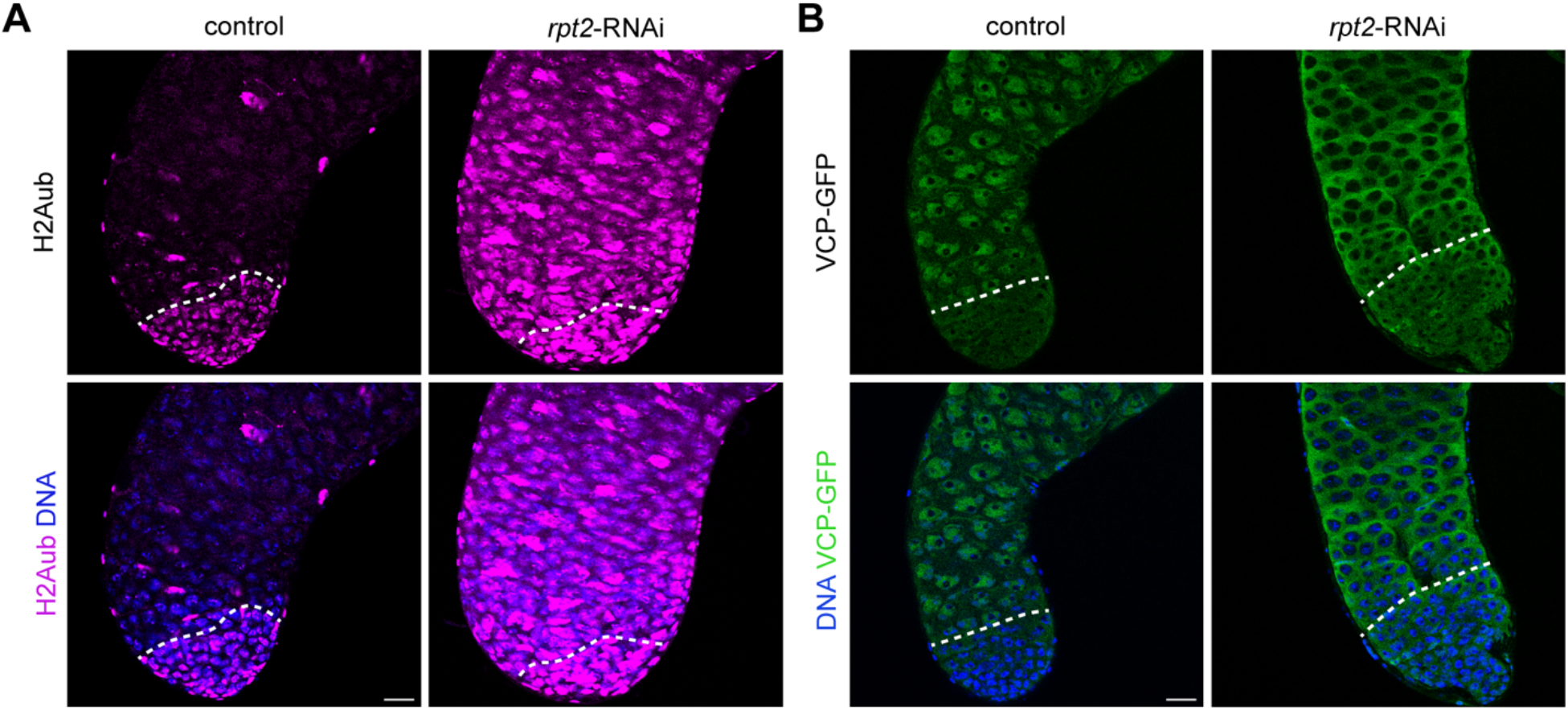
The proteasome component, Rpt2, is required for H2Aub downregulation and VCP nuclear entry. (**A**) Images of H2Aub and Hoechst (DNA) in control (BamGal/+) and *rpt2*-RNAi (BamGal>*rpt2*-RNAi) testes. The dashed lines indicate the mitotic-meiotic transition. (**B**) Images of VCP-GFP and Hoechst (DNA) in control and *rpt2*-RNAi (BamGal>*rpt2*-RNAi) testes. The dashed lines indicate the mitotic-meiotic transition. Bars, 20 μm.

**Figure S7.**
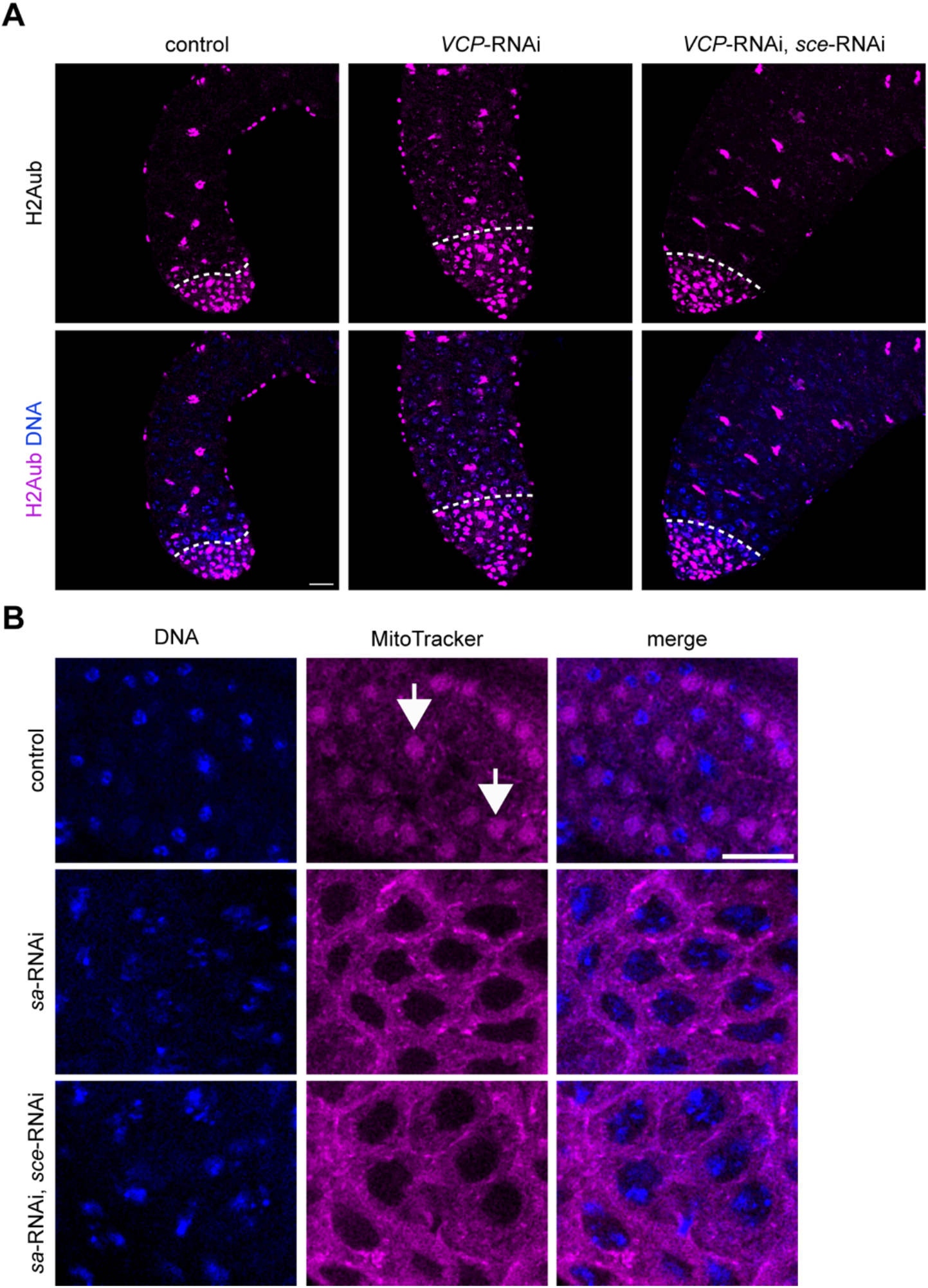
Knockdown of *sce* blocks H2Aub but is insufficient to promote spermatocyte differentiation in the absence of Sa. (**A**) Images of H2Aub and Hoechst (DNA) in control (BamGal/+), *VCP*-RNAi (BamGal>*VCP*-RNAi), and DKD (BamGal>*VCP*-RNAi, *sce*-RNAi) testes. The dashed lines indicate the mitotic-meiotic transition. (**B**) Images of MitoTracker and Hoechst (DNA) in control (BamGal/+), *sa-* RNAi (BamGal>*VCP*-RNAi), and *sa*-RNAi/*sce*-RNAi (BamGal>*sa*-RNAi, *sce*-RNAi) testes. The most developed germ cells in *sa*-RNAi and *sa*-RNAi/*sce*-RNAi testes were spermatocytes (note the presence of bivalents and the absence of a nebenkern). Round spermatids are shown in control testes for comparison (note the compact chromatin morphology and the nebenkern [arrows]). Bars, 20 μm.

**Figure S8.**
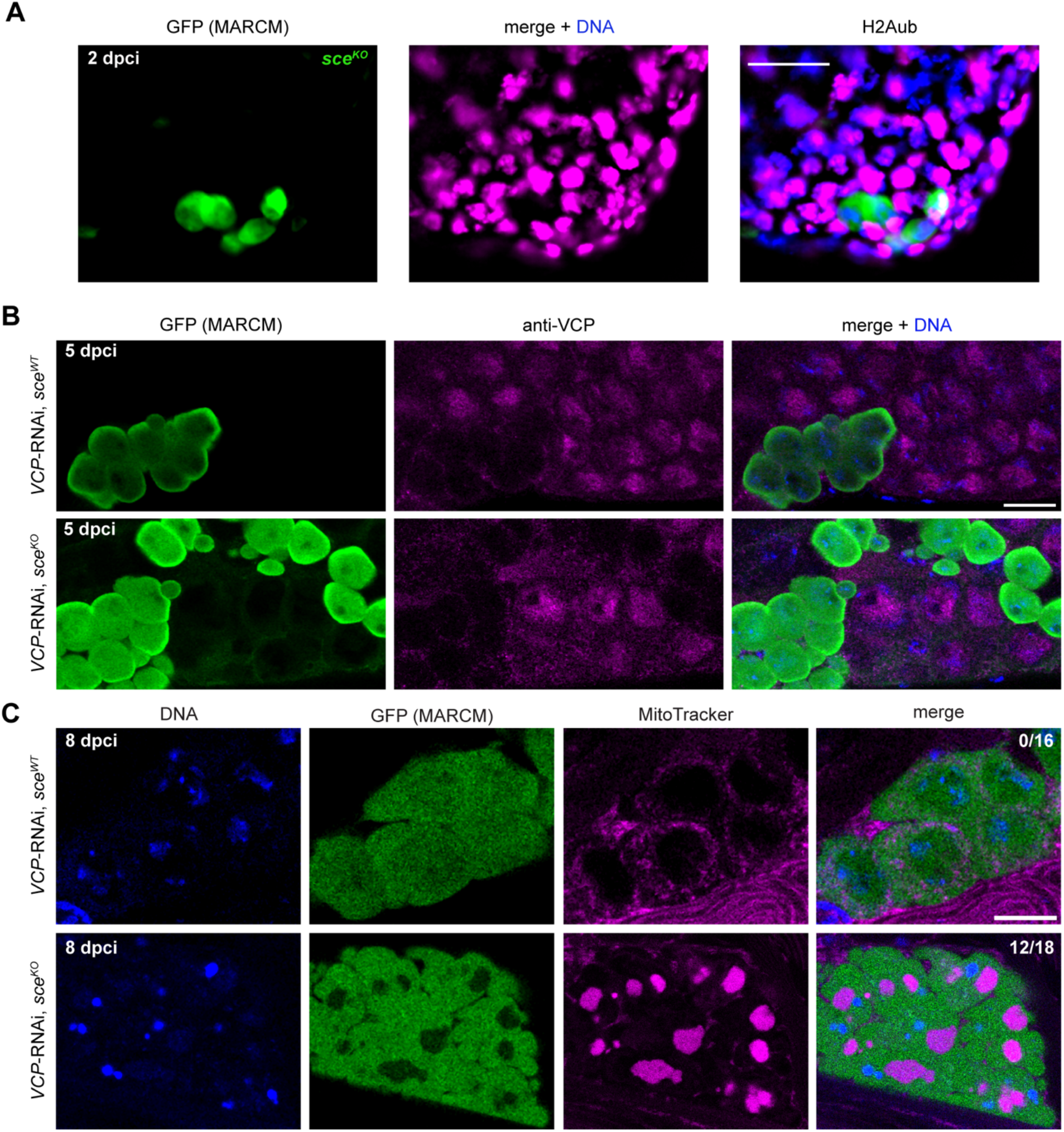
Suppression of H2A mono-ubiquitination supports spermatocyte differentiation in the absence of VCP. (A) Images of GFP (*sce^KO^* MARCM clones), H2Aub, and Hoechst (DNA) in spermatogonia at 2 dpci. (B) Images of GFP (MARCM clones), VCP (anti-VCP), and Hoechst (DNA) in spermatocytes at 5 dpci. In the *VCP*-RNAi, *sce^WT^* panel, GFP-positive MARCM clones express *VCP*-RNAi and are *sce^WT^* homozygotes. In the *VCP*-RNAi, *sce^KO^* panel, GFP-positive MARCM clones express *VCP*-RNAi and are *sce^KO^* homozygotes. (C) Images of GFP (MARCM clones), Hoechst (DNA), and MitoTracker in testes at 8 dpci. In the *VCP*-RNAi, *sce^WT^* panel, GFP-positive MARCM clones express *VCP*-RNAi and are *sce^WT^* homozygotes; these cells are spermatocytes, as indicated by the large cell size, dim Hoechst labeling, separation of bivalents, and small, punctate mitochondria. In the *VCP*-RNAi, *sce^KO^* panel, GFP-positive MARCM clones express *VCP*-RNAi and are *sce^KO^* homozygotes; these cells are round spermatids, indicated by the small cell size, bright Hoechst staining, compacted chromatin, and large, clustered mitochondria adjacent to nuclei. Numbers in the right-most images indicate the proportion of testes containing post-meiotic germ cells. Bars, 20 μm.

